# Cryogenic volume electron microscopy of whole plant protoplasts

**DOI:** 10.64898/2025.12.15.694529

**Authors:** Xiao Zhang, Zihan Lin, Xinlin Chen, Jake Kaminsky, Liguo Wang, Qun Liu

**Affiliations:** Biology Department, Brookhaven National Laboratory, Upton, NY 11973, USA; LBMS, Brookhaven National Laboratory, Upton, NY 11973, USA; NSLS-II, Brookhaven National Laboratory, Upton, NY 11973, USA

## Abstract

Volume electron microscopy (vEM) provides nanometer-scale, three-dimensional imaging of cells, but applying it to plant systems remains challenging. Cell walls, large vacuoles, and tissue thickness complicate sample preparation and cryogenic imaging. Here we report a cryogenic vEM (cryo-vEM) workflow for unstained plant protoplasts that achieves volumetric imaging of whole vitrified sorghum stem protoplasts without chemical fixation, dehydration, resin embedding, or heavy-metal staining. The method integrates optimized protoplast isolation, plunge-freezing vitrification for native-state preservation, automated cryogenic focus ion beam scanning electron microscopy (cryo-FIB-SEM) slice-and-view acquisition, contrast enhancement and stack alignment, and AI-assisted human-in-the-loop 3D segmentation. Using sorghum stem protoplasts as a demonstration, the workflow captured large-volume frozen-hydrated protoplast ultrastructure, allowing visualization of major organelles, including the nucleus, mitochondria, vacuoles, ER/Golgi-like membranes, lipid bodies, and subcellular features consistent with nuclear-envelope pores. We further quantified organelle volumes and surface areas from the segmented 3D data, highlighting the potential for quantitative cellular ultrastructure analysis. This cryo-vEM workflow provides a platform for near-native structural studies of isolated plant protoplasts.

## Introduction

Volumetric cellular ultrastructure imaging using focused ion beam scanning electron microscopy (FIB-SEM) has been revolutionizing cell biology. Volume electron microscopy (vEM) has emerged as a powerful technology to achieve nanometer-resolution 3D imaging of whole cells ^1-5^. Techniques like serial block-face SEM and FIB-SEM allow researchers to iteratively section a sample and image each newly exposed layer with an electron beam, reconstructing a high-resolution volume. Recently, FIB-SEM has demonstrated near isotropic resolutions well below 10 nm in all three dimensions ^1,6^. This capability has made vEM attractive in studying mammalian cell biology for mapping organelle architectures and interactions in 3D, from endoplasmic reticulum–mitochondria contact sites to neuronal synapses ^1,3,6-8^. The past decade has seen vEM applied extensively to animal and microbial cells, producing detailed atlases of cellular organization and ultrastructural changes of organelles across cell types and development stages. These studies highlight an emerging role of vEM connecting 3D ultrastructure to function.

Applying vEM to plant systems remains less routine than in many animal or microbial samples. The cell wall is not a universal barrier to fixation or staining, but plant tissues often combine rigid walls, large vacuoles, extracellular spaces, heterogeneous cell types, sample thickness, and variable reagent penetration. These features can make conventional fixation/staining and cryogenic workflows sample-dependent and can contribute to uneven contrast, charging, or incomplete vitrification. As a result, many plant vEM studies have relied on specialized fixation, dehydration, embedding, and heavy-metal staining protocols to generate sufficient contrast for organelle-scale imaging ^9,10^.

Cryogenic preservation offers an alternative way of electron microscopy (EM) sample preparation. By rapidly freezing hydrated cells, cryogenic workflows can preserve cellular structures close to their native hydrated state and avoid chemical fixation, dehydration, heavy-metal staining, and resin embedding, which may introduce structural artifacts. However, cryogenic plant imaging introduces its own constraints, including limited plunge-freezing depth, low intrinsic contrast, and the need to thin thick samples before high-resolution cryogenic imaging ^7,11,12^.

Cryo-electron tomography (cryo-ET) and cryogenic volume electron microscopy (cryo-vEM/cryo-FIB-SEM) are complementary cryogenic EM techniques. Cryo-ET enables molecular-scale visualization within small volumes, whereas cryo-vEM reconstructs substantially larger volumes at organelle-scale resolution. Recent plant cryo-ET studies have imaged Arabidopsis root protoplasts, isolated/intact chloroplasts, and intact plant tissues or specialized cell wall regions by combining high-pressure freezing, cryo-CLEM targeting, cryo-FIB milling/lift-out, and tomography ^13-16^. These studies provide high-resolution structural information in near-native contexts, but cryo-ET is inherently restricted to small lamella volumes and often requires technically demanding sample preparation, high-pressure freezing, cryo-CLEM targeting, and cryogenic lift-out.

Cryo-FIB-SEM volume imaging has also recently been demonstrated in *Vicia faba* guard cells, enabling three-dimensional reconstruction of organelles in vitrified higher-plant tissues ^17^. This study established the feasibility of cryogenic vEM imaging in higher plants but focused on intact epidermal guard cells and did not address other cell types or quantitative analysis of whole-cell ultrastructure. Consequently, a workflow that combines large-volume cryogenic imaging with quantitative analysis of plant protoplast ultrastructure remains lacking.

Plant protoplasts are cells with cell walls removed by digesting enzymes and have been widely used in plant biology ^14,18,19^. They retain the plasma membrane and many subcellular organelles, making them useful for cellular and molecular studies. Protoplasts are not equivalent to intact walled cells because cell-wall removal can alter cell shape, cytoskeletal organization, vacuolar architecture, and organelle positioning. However, their reduced thickness and accessibility nevertheless make them useful for developing cryo-vEM imaging and segmentation workflows.

Here we present a cryo-vEM workflow for imaging whole plant protoplasts that leverages plant protoplast preparation, protoplast vitrification, cryo-FIB-SEM data acquisition, and computational tools for imaging processing and segmentation. The workflow produces organelle-scale 3D volumes of whole frozen-hydrated protoplasts with lateral pixel sizes down to 4 nm under selected acquisition settings. The workflow provides a platform for volumetric imaging and quantification of plant protoplasts.

## Results

### Development of a cryo-vEM workflow for plant protoplasts

To address the challenges of applying vEM for plant protoplasts, we developed an end-to-end workflow for plant protoplasts (**Fig. 1**). Sorghum stem tissues were enzymatically digested to isolate protoplasts, which were then deposited onto poly-L-lysine-treated cryo-EM grids to improve cell adhesion during sample preparation. Following plunge freezing in liquid ethane, vitrified protoplasts were imaged using automated cryo-FIB-SEM slice-and-view data acquisition to generate volumetric datasets. The acquired image stacks were subsequently processed through contrast enhancement and stack alignment to improve image quality and inter-slice continuity. Finally, because plant cryo-vEM datasets remain scarce, we employed a human-in-the-loop segmentation workflow to iteratively generate and refine 3D segmentation masks (**Supplementary Fig. 1**), enabling quantitative analysis of organelle morphology and cellular organization.

**Figure 1.**
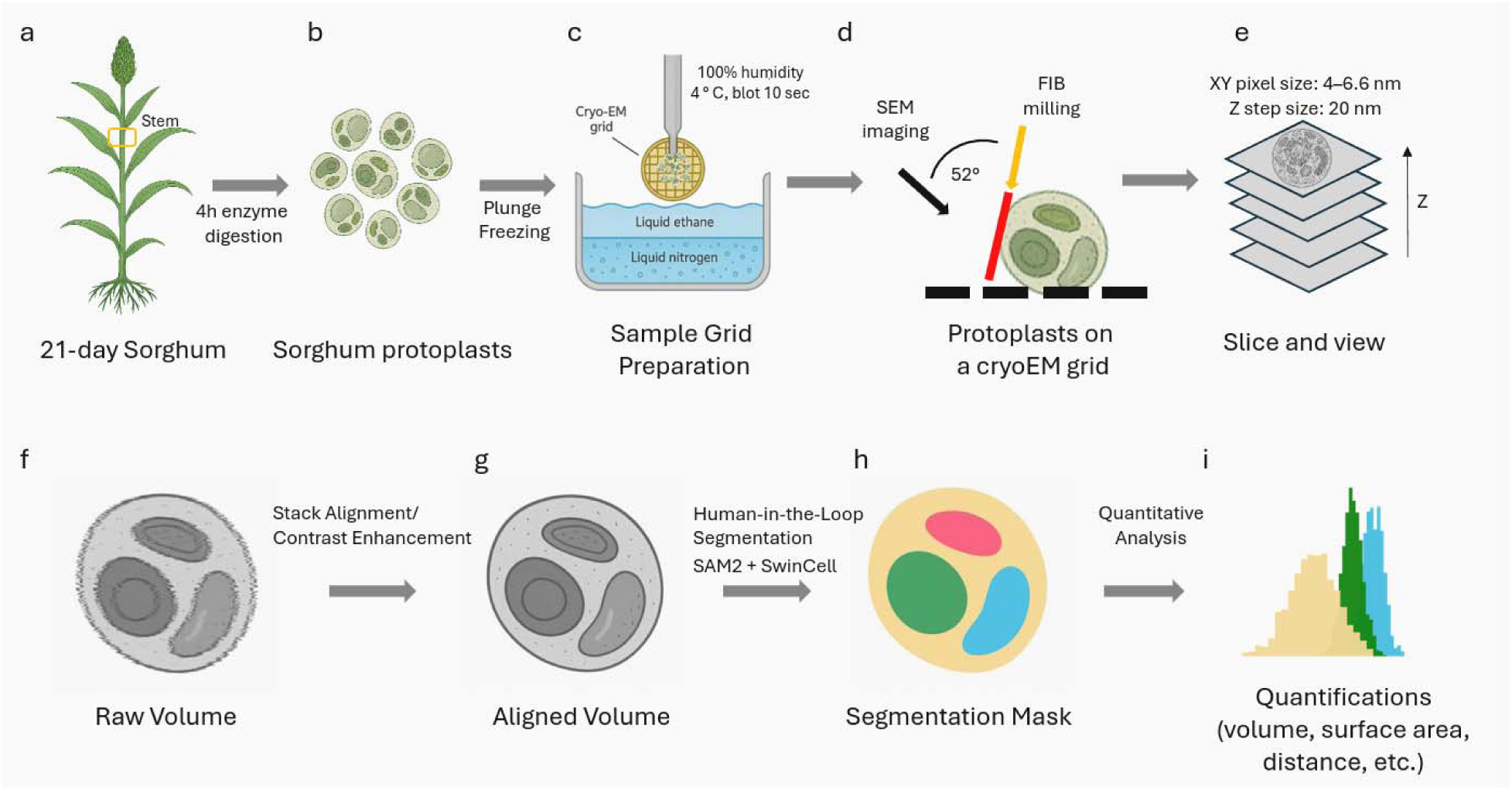
Cryo-VEM workflow for plant protoplasts. (a,b) Isolation of protoplasts from sorghum stem tissue. Stem tissues from 21-day-old sorghum seedlings were enzymatically digested for 4 h to release protoplasts. (c) Cryo-vitrification of protoplasts on cryo-EM grids. Protoplasts were deposited onto poly-L-lysine-treated cryo-EM grids and plunge-frozen in a Vitrobot chamber maintained at 4 °C and 100% humidity following 10 s of manual backside blotting. (d) Serial cryo-FIB milling and SEM imaging of vitrified protoplasts using a slice-and-view workflow. (e) Representative SEM image stack acquired from the cryo-FIB-SEM experiment with lateral pixel sizes of 4–6.6 nm and a slice spacing of 20 nm in the z direction. (f) Reconstructed raw 3D volume generated from the image stack. (g) Post-processed volume following CLAHE-based contrast enhancement and SIFT-based stack alignment. (h) AI-assisted, human-in-the-loop segmentation of subcellular structures using SAM2 and SwinCell. (i) Quantitative analysis of segmented organelles, including measurements of organelle volume, surface area, and spatial organization.

### Cellular ultrastructure by cryo-vEM

Protoplasts lack the structural support of cell walls. They are mechanically fragile and highly sensitive to osmotic and shear stress. To minimize damage to cells, we extracted protoplasts gently and at low shear forces and deposited them on cryoEM grids with optimized buffer conditions. The workflow yielded large-volume cryo-FIB-SEM datasets from multiple sorghum stem protoplasts. These datasets allowed organelle-level 3D visualization, although image quality was not uniform across all regions because local charging, curtaining, and focus variation reduced interpretability in parts of some volumes.

**Fig. 2a** and **b** show orthogonal views of a reconstructed 3D volume acquired from a vitrified sorghum protoplast (**Supplementary Movie 1**). Representative slices from the XY and XZ planes illustrate the overall cellular morphology. Most intracellular organelles, including nucleus (Nuc), mitochondria (Mito), lipid droplets (LBs), vacuole (VA), and endoplasmic reticulum (ER)/Golgi-like membranes can be visualized. Small vesicle-like particles are also visible, but their identity cannot be assigned confidently from the present contrast and resolution. Sorghum stem tissues contain heterogeneous cell populations with distinct physiological functions and organelle compositions. Consistent with this heterogeneity, the protoplast datasets analyzed in this study showed variation in plastid content. In the representative protoplast shown in **Fig. 2a**, plastids were not readily identifiable, whereas additional protoplast datasets contained amyloplast-like organelles with prominent starch granules (**Supplementary Fig. 2**). These observations indicate that plastid occurrence varies among sorghum stem-derived cell populations.

**Figure 2.**
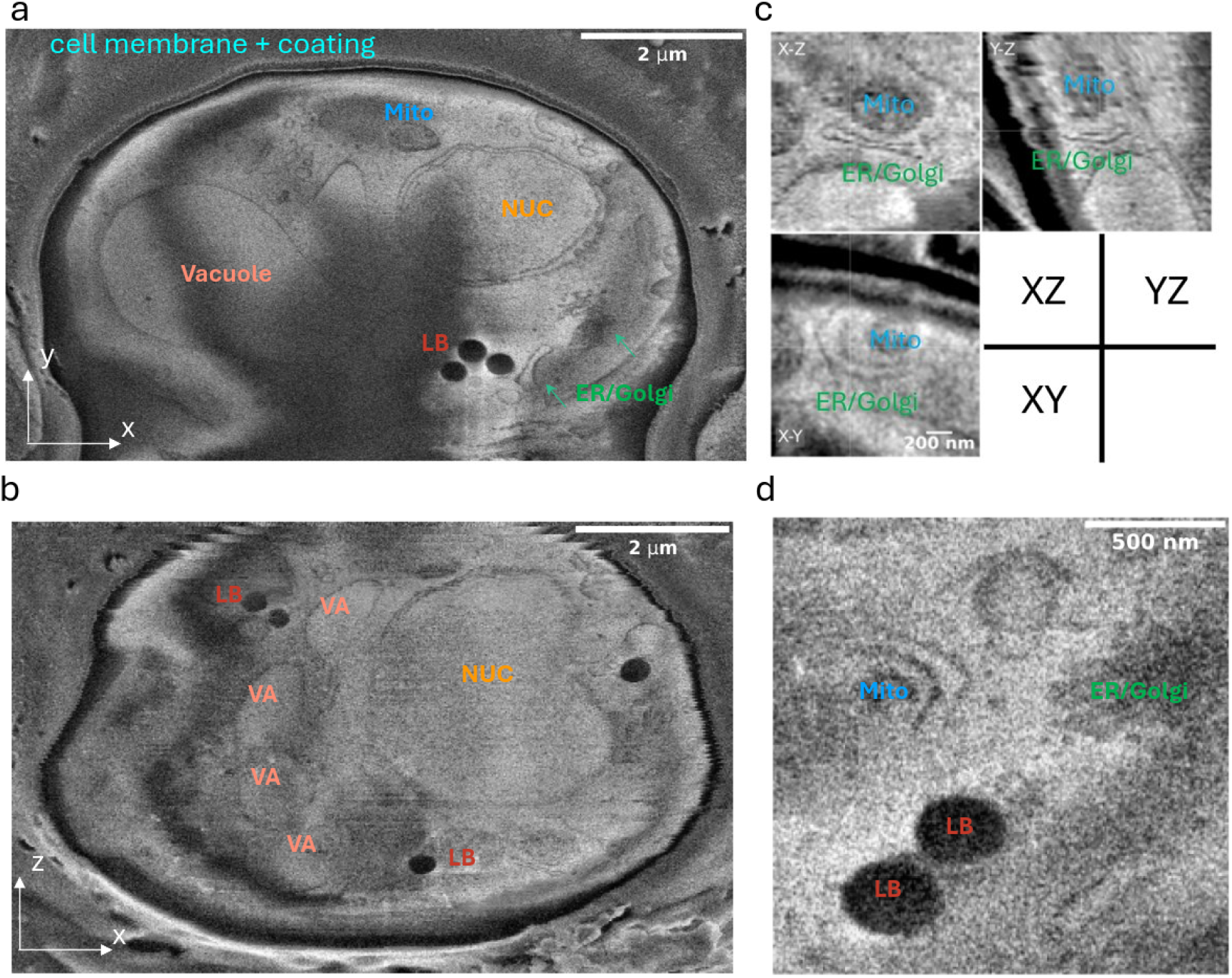
Ultrastructure of a sorghum protoplast by cryo-VEM. (a,b) Representative views from a reconstructed 3D volume EM of vitrified sorghum protoplast. (c) three orthogonal views of one region of interest (ROI) containing a region showing close spatial association between ER/Golgi-like membranes (green) and a mitochondrion (cyan). (d) Another ROI from the 3D volume showing dense arrangement of lipid bodies (LB), ER/Golgi-like membranes, and mitochondria.

**Fig. 2c** shows three orthogonal views of one region of interest (ROI), containing a curved membrane structure closely associated with a mitochondrion. This morphology suggests close organelle proximity, but the current cryo-vEM data do not allow definitive identification of a bona fide membrane contact site or assignment of the membrane identity. Higher-resolution cryo-ET or molecular marker labeling would be required to confirm membrane identity and contact-site architecture. Another ROI illustrates the dense organelle architecture in the cytoplasm, including spatially close arrangements of lipid bodies (LB), ER/Golgi-like membrane structures, and mitochondria (**Fig. 2d**).

### Post-processing of cryo-vEM images

To preserve near-native state ultrastructure, no chemical fixation or heavy metal staining was used in the sample preparation steps. While this approach avoids fixation and staining artifacts, it may result in relatively low image contrast. To improve interpretability, we implemented a digital post-processing pipeline that includes local contrast enhancement using contrast-limited adaptive histogram equalization (CLAHE) ^20^, a well-established method. **Supplementary Fig. 3a,b** shows a representative 2D slice of the cell before and after local contrast enhancement by CLAHE. Quantitative analysis of Shannon entropy across all slices (**Supplementary Fig. 3e**) showed a significant increase in image information content with CLAHE.

The vEM workflow acquires 2D slices sequentially and can take many hours, making it sensitive to temperature, mechanical, or charging-induced drifts. These drifts may introduce misalignment and make the 3D image analysis challenging. To enhance 3D image quality, the slices of a volume were aligned using Scale-Invariant Feature Transform (SIFT) registration ^21^. Feature points and descriptors are extracted from consecutive slices and then aligned. **Supplementary Fig. 3c,d** shows the side view of one ROI, including membrane structures before and after SIFT alignment. Before the SIFT alignment, membrane contours appear jagged and discontinuous. After the SIFT alignment, the membrane structure and vesicle are clearly resolvable. The estimated slice-to-slice drifts in the x and y directions are plotted in **Supplementary Fig. 3f**. CLAHE and SIFT are established methods used here as practical components of the workflow.

After these post-processing steps, the cryo-vEM workflow yielded contrast-enhanced volumetric images of subcellular ultrastructure for organelle-level interpretation. **Fig. 3a** presents four XY slices extracted from a single ROI at different Z positions. Notably, features consistent with nuclear-envelope pores, which usually have sizes around 50 nm, are visible in the second and third slices. Nuclear-envelope pores are essential for selective nucleocytoplasmic transport of macromolecules and play key roles in regulating gene expression and signaling ^22^. Revealing its position and structure may help in studying communication and interactions between nuclear and other organelles. In addition to nuclear pores, we observed a cluster of a group of vesicles localized between two large vacuoles (**Fig. 3b**). Vacuoles of plant cells are essential for dynamic processes of storage, ion balance, and stress response. The localization of these vesicles between vacuoles indicates close spatial association between these compartments. The identity and functional significance of these vesicles cannot be determined from the current dataset and will require higher-resolution imaging and molecular markers. Collectively, we demonstrate that our cryo-vEM workflow can support organelle-level interpretation of protoplast ultrastructure, while higher-resolution imaging will be needed for molecular assignment.

**Figure 3.**
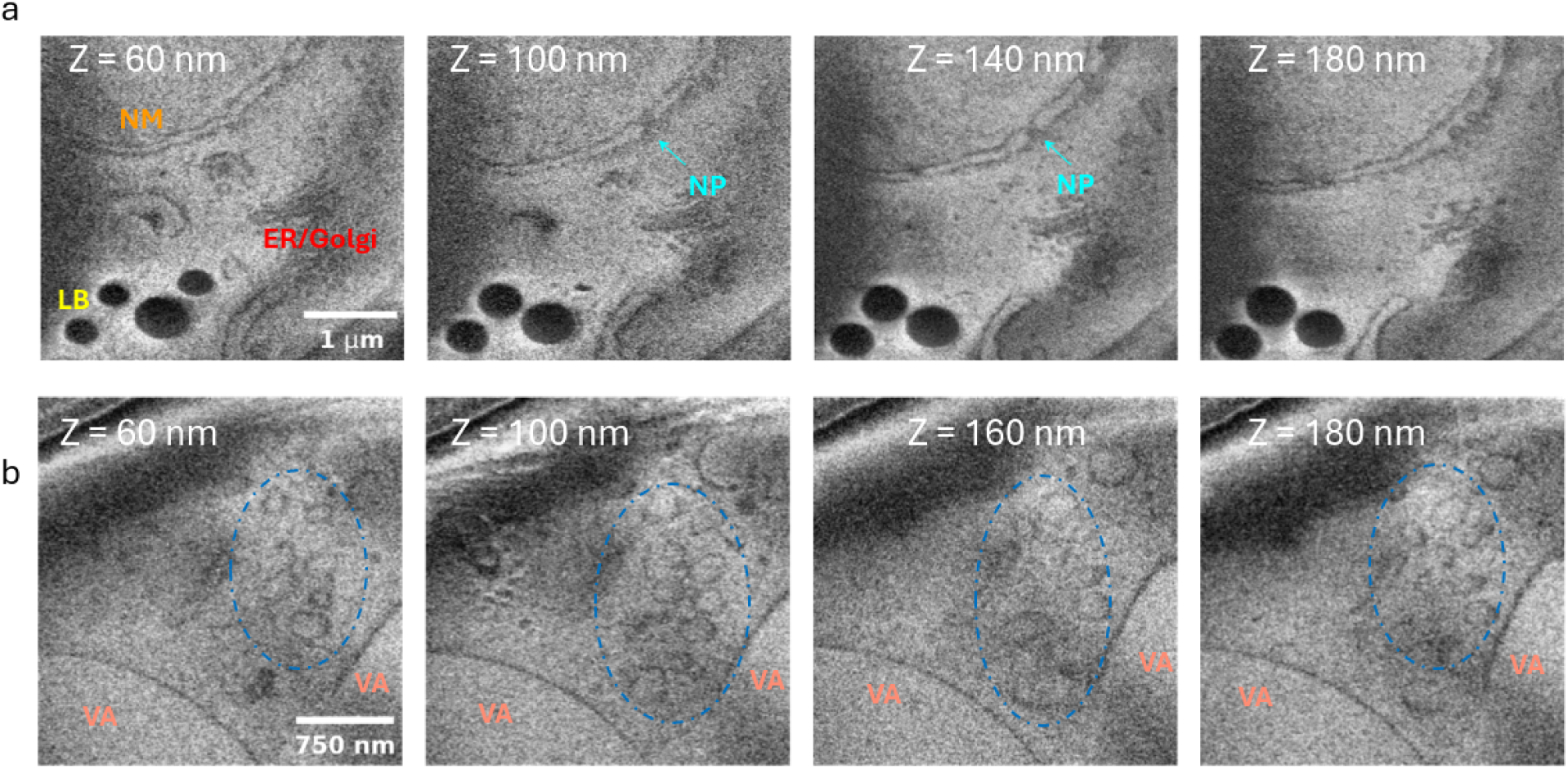
Representative slices extracted from the cryo-fixed sorghum protoplast volume. (a) Four slices from different z positions showing key subcellular features. Nuclear membrane (NM, Orange), Nuclear-pore-like feature (NP, arrowhead), ER/Golgi membranes, Lipid droplets (LB), (b) Four slices acquired at different z positions showing a cluster of vesicle-like structures (blue dashed ovals) located between two vacuoles (VA). The morphology and spatial organization of these structures are visible across multiple consecutive slices, demonstrating the continuity of membrane-rich features in the reconstructed volume.

### AI-assisted human-in-the-loop segmentation and quantification

After the post-processing steps, we performed human-in-the-loop segmentation of organelles observed in the vEM dataset using SAM2 ^23^ and SwinCell ^24^ to produce 3D segmentation masks. We then performed 3D volume rendering and quantitative analysis of the segmented organelle classes of nucleus, mitochondria, vacuole, lipid droplet, ER/Golgi-like membranes, and plasma membrane (**Fig. 4a**). The segmented masks were used to compute both total volume and surface area for each segmentation class, providing an integrated view of organelle size and morphology. Among these segmented classes, the nucleus occupied the largest volume, followed by vacuoles, reflecting their well-preserved native like states during sample preparation and volumetric imaging. Our segmentation and cellular analysis provide quantitative measurements for future studies of organelle remodeling in response to environmental stress or pathogen interactions.

**Figure 4.**
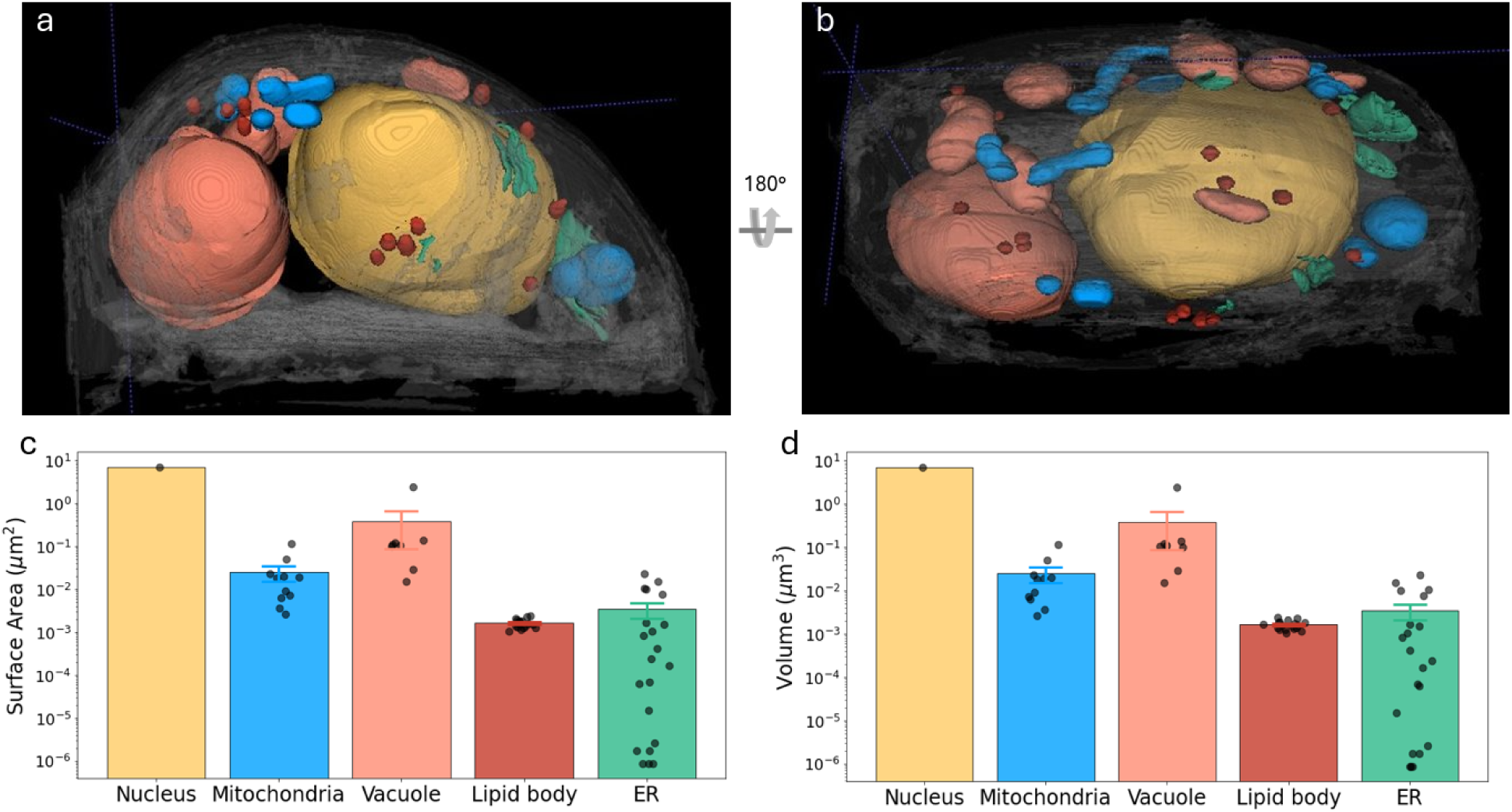
3D volume rendering and quantification of segmented protoplast organelles. (a,b) Two views of the segmented volume. (c) Calculated surface area of the segmented organelles. (d). Calculated volume of the segmented organelles.

## Discussion

In this study, we developed a cryogenic volume electron microscopy (cryo-vEM) workflow for organelle-scale 3D imaging of plant protoplasts. Our workflow integrates optimized protoplast preparation, plunge-freezing vitrification, automated cryoFIB-SEM acquisition, post-processing, and AI-assisted human-in-the-loop segmentation. This method enables visualization and quantification of major subcellular ultrastructure in unstained, frozen-hydrated sorghum stem protoplasts without chemical fixation or heavy metal staining being used. The workflow demonstrates the applicability of cryo-vEM to study plant protoplasts through a structural approach, enabling visualization and quantification of cellular organelles.

Compared to conventional vEM workflows, which rely on dehydration, resin embedding, and staining steps that can distort membrane morphology and organelle integrity, our approach preserves protoplasts in a frozen-hydrated state, avoiding processing steps that can distort membrane morphology. The use of protoplasts removes the wall and simplifies grid deposition, sample thickness, plunge freezing, and cryo-FIB-SEM access for this workflow. However, several limitations should be noted. First, wall removal can alter cell shape, vacuolar organization, cytoskeletal organization, and organelle positioning. Thus, conclusions from this workflow should be limited to protoplasts. Second, large-volume cryo-vEM datasets remain susceptible to charging, curtaining, and focus-related artifacts, which can reduce image quality and obscure ultrastructural features in some regions of the reconstructed volume. These limitations also affect downstream segmentation accuracy, particularly for organelles with complex membrane morphology or ambiguous ultrastructural appearance. Nevertheless, the current workflow enables quantitative analysis of major organelle classes. Future improvements in image quality, together with the development of larger annotated plant cryo-vEM datasets and plant-specific deep-learning models, are expected to further improve segmentation performance and organelle classification. Third, because the workflow is based on protoplasts, future extension of the workflow to intact tissues will be needed to improve robustness and biological applicability.

A particular challenge in preserving plant cells for imaging is the presence of a large number of vacuoles of various sizes and shapes (**Fig. 2**), which are highly susceptible to collapse during chemical fixation and staining. In our workflow, vitrification process appeared to preserve vacuolar shapes and maintained their positioning in cytoplasm, enabling the visualization of both intact vacuolar morphology and their spatial arrangement relative to adjacent organelles, underscoring the value of this method for capturing native-state architecture in plant protoplasts.

Segmenting large volume EM data is labor-intensive, particularly for complex biological structures with densely packed organelles. Although deep learning-based segmentation models have shown success in mammalian datasets ^6,25^, these models are generally trained on chemically fixed and heavy-metal-stained samples and may not be applicable directly to cryogenic volumetric data. In addition, volume EM datasets of plant cells are still rare; existing models do not generalize well to plant cells due to significant differences in subcellular contents and morphologies.

Among subcellular structures, segmentation of the nucleus was relatively straightforward because of its size, shape, and contrast. SAM2 ^23^ was used as a generalist foundation segmentation model pretrained by its developers on large-scale natural image and video datasets rather than on plant or electron microscopy data. We used SAM2 interactively to generate initial masks for the nucleus and cytosolic regions. More complex organelles, including ER/Golgi-like membranes and mitochondria, required manual curation and iterative SwinCell ^24^ training because pretrained generalist models did not provide reliable masks for low-contrast cryogenic plant vEM data. This human-in-the-loop approach reduced manual annotation burden, but larger plant cryo-vEM training datasets and improved acquisition quality will be needed before robust automated segmentation is possible.

The overall workflow can be completed within a practical timeframe. Protoplast isolation typically requires 6-8 h. Grid preparation consists of poly-L-lysine (PLL) treatment for approximately 1 h, followed by overnight drying under vacuum. Once prepared, sample deposition, manual blotting, and plunge freezing require approximately 2 h. Cryo-FIB-SEM acquisition of a single volume requires approximately 20 h under the imaging conditions used in this study. The subsequent image-processing pipeline, including CLAHE-based contrast enhancement, SIFT-based stack alignment, and AI-assisted segmentation, can be completed within a week depending on the extent of manual curation required. Although human-in-the-loop segmentation remains the most time-consuming computational step, it reduces the annotation burden relative to fully manual annotation. Collectively, these processing times show that the workflow is feasible for plant protoplast cryo-vEM experiments, while further automation and improved imaging quality will be needed for higher-throughput applications.

Our workflow complements recent advances in plant cryogenic electron microscopy. Franzisky et al. ^17^ demonstrated the feasibility of cryo-FIB-SEM volume imaging in Vicia faba guard cells and established an important foundation for applying cryogenic volume imaging to higher-plant tissues. Similarly, Daraspe et al. ^16^ and Pöge et al ^15^ developed cryo-CLEM, high-pressure-freezing, cryo-FIB milling/lift-out, and cryo-ET strategies for targeted imaging of intact plant tissues and specialized cell wall regions. Wietrzynski et al ^13^ further demonstrated the power of cryo-ET for visualizing molecular architecture in intact spinach chloroplasts. In comparison, our workflow focuses on isolated plant protoplasts and integrates cryo-vEM with human-in-the-loop segmentation and quantitative organelle analysis. We view cryo-vEM and cryo-ET as complementary: cryo-vEM provides larger-volume cellular context, whereas cryo-ET provides higher-resolution structural information in selected regions of interest.

## Methods

### Protoplast isolation

Protoplasts were isolated from sorghum stems using a protocol adapted from reference ^18^. Twenty-one-day-old seedlings were harvested and leaves along with the upper stem section were discarded. The remaining stems were rinsed thoroughly with Milli-Q water and gently dried with Kimwipes. Stem and mid-vein tissues were cut into ∼1 mm segments and incubated in 10 mL of freshly prepared, filter-sterilized enzyme buffer (0.5 M mannitol, 10 mM KCl, 1 mM CaCl_₂_, 8 mM MES pH 5.7, 0.6% cellulase from *Aspergillus niger* (Sigma-Aldrich, 22178-25G), 0.1% pectinase from Rhizopus sp. (Sigma-Aldrich, P2401-5KU), 0.4% polyvinylpyrrolidone K15). Digestion proceeded for 4 h at 27°C and 30 rpm in the dark. Subsequently, 10 mL of W5 solution (154 mM NaCl, 125 mM CaCl_₂_, 5 mM KCl, 2 mM MES, pH 5.7) was added, and incubation continued for 1 h under the same conditions. The suspension was filtered through a 70 μm cell strainer (Falcon, 08-771-2) into a 50 mL tube on ice. All subsequent steps were performed on ice. The filtrate was centrifuged at 200 × g for 5 min at 4°C in a swing-bucket rotor. The supernatant was carefully removed to eliminate residual enzyme solution. The pellet was gently resuspended in at least 1 mL of W5 solution and recentrifuged (200 × g, 5 min, 4°C). The supernatant was discarded, and the pellet resuspended in 1 mL of W5 solution. This buffer exchange was repeated four times. Protoplast concentration was determined by manual counting using a hemocytometer (**Supplementary Fig. 4a**), yielding approximately 5 × 10□ protoplasts/mL.

### CryoEM grid and cryo-VEM sample preparation

Grids were pretreated to enhance protoplast adhesion. Quantifoil R 3/3, 200 mesh Cu grids (Quantifoil Q250CR3) were glow-discharged using a PELCO easiGlow Glow Discharge Cleaning System (Ted Pella, Inc.) at 15 mA for 90 s, treating the back side first and then the front (carbon) side. Glow-discharged grids were transferred to a clean glass slide with the carbon side facing up. A 3 μL aliquot of 0.5% poly-L-lysine (Sigma-Aldrich, P4707) was applied to each grid, and the slide was placed in a Petri dish for incubation at 37°C for 10 min. To prevent drying, 50 μL of autoclaved Milli-Q water was added to each grid. Grids were then lifted with forceps and blotted from the back side using filter paper. Next, 6 μL of autoclaved Milli-Q water was applied to the carbon side. After 10 s, excess water was blotted from the back side with filter paper. This washing step was repeated once. Finally, grids were stored in a Petri dish under vacuum overnight upon use.

Cryo-EM grid preparation was optimized by screening grid type, pretreatment, protoplast concentration, blotting temperature, and ice thickness, modifying protocol ^14^. Initially, equal volumes of protoplasts were applied to glow-discharged continuous carbon grids under varying conditions. Pretreatment with 0.5% poly-L-lysine yielded the highest protoplast adhesion after manual blotting. Excess protoplasts resulted in excessively thick ice after plunge freezing. Optimal conditions were determined to be 3 μL of 5 × 10□ protoplasts/mL, which produced a single layer with distinguishable cell-to-cell spacing. Ice thickness was assessed visually on the AutoGrid loading stage (**Supplementary Fig. 4b,c**) and further verified by cryo-TEM imaging (**Supplementary Fig. 5b,c**) to ensure suitability for subsequent cryo-vEM experiments.

After screening and optimization of grid and sample conditions, 4 μL of 5 × 10□ protoplasts/mL was applied to the pre-treated grids. Vitrobot was set to 4°C and 100% humidity without automated blotting. Grids were blotted manually from the back for 10 s, followed by plunge-freezing immediately. Frozen grids were stored in liquid nitrogen for cryo-VEM analysis.

### Volume Electron Microscopy Data Acquisition

Volume EM imaging was performed using an Aquilos 2 DualBeam cryo-FIB-SEM system (Thermo Scientific). Whole-grid atlas images were acquired at low magnification (50×) using the MAPS software to identify well-preserved protoplasts. The sample surface was sputter-coated at 10 Pa, 30 mA for 10 s, then Pt GIS deposition was applied for 30 seconds to reduce certain artifacts. Then the sample surface was sputter-coated again with 10 Pa, 30 mA for 10 s.

After coating, the region of interest (ROI) was centered, and a cross-shaped fiducial marker was milled near the target area using a 0.5 nA gallium ion beam to aid beam alignment and navigation during automated slice-and-view milling/imaging processes.

To expose the internal cell structures for imaging, a front trench was milled in front of the target protoplast using a 0.5 nA ion beam. A left-side trench was also milled at the same current to mitigate sample charging during imaging. Serial sectioning and imaging (slice-and-view) were performed using a focused ion beam set to 50 pA for fine milling, with a milling step size of 20 nm. The acquired image resolution was 1958×1743 pixels, resulting in a pixel size of 6.6 nm. For selected regions of interest, a subset of slices was collected with 3230×2877 resolution, resulting in a pixel size of 4 nm.

Images were acquired using the Everhart-Thornley Detector (ETD) with an electron beam acceleration voltage of 30 keV, dwell time 100 ns, and 64-line integration with no frame integration. The chamber temperature was maintained below –170°C throughout data acquisition to preserve the vitrified state of the sample.

### Cryo-vEM data processing

Local contrast in raw cryo-vEM images was enhanced using Contrast Limited Adaptive Histogram Equalization (CLAHE) ^20^. An ImageJ macro script was developed to apply this algorithm to each 2D slice. CLAHE enhances fine features such as membrane boundaries and vesicular structures while suppressing low-frequency background noise. The macro script was configured with a block size of 127 and 256 histogram bins, with input parameters being optimized to enhance cellular visibility without introducing artifacts. This step significantly improved membrane continuity and organelle boundary clarity across the stack.

To correct for drift and misalignment between adjacent 2D slices, we performed affine alignment using OpenCV’s ^26^ implementation of Scale-Invariant Feature Transform (SIFT) ^21^. First key points and corresponding descriptors were extracted using OpenCV’s ‘*detectAndCompute’* function, and then the extracted key points were matched using the ‘*BFMatcher’* function. A transformation matrix was calculated based on the matched key points, then warpAffine was applied to align consecutive slices. The same operations were performed between consecutive slices, with each slice registered to the previous one, and eventually, a 3D volume. This method enables 3D image data analysis by improving inter-slice continuity and enables accurate 3D reconstruction, particularly enhancing the interpretability of fine-detail regions.

### vEM image segmentation

Six distinct subcellular components were segmented from the volume EM datasets: nucleus, mitochondria, ER/Golgi-like membranes, lipid droplets (LBs), vacuole, and plasma membrane. Segmentation was performed using a human-in-the-loop deep learning approach with manual curation to ensure both accuracy and scalability.

For relatively regular high contrast structures such as the plasma membrane, nucleus, and lipid droplets, initial masks were generated with the interactive segmentation tool SAM2 ^23^. One representative 2D slice from the 3D volume was annotated with positive (foreground) and negative (background) point prompts (**Supplementary Fig. 1b**). Then, the SAM2 model used the prompts to propagate segmentation masks to other 2D slices (**Supplementary Fig. 1c**). The segmented 2D slices were then stitched together to get 3D masks.

For organelles with more complex and irregular morphologies, such as the ER, we used a semi-automated workflow. Initial masks were manually generated with SAM2 and used as training data for a Swin-Transformer-based SwinCell model. The model was trained to predict organelle masks on unannotated slices, and predictions were iteratively refined. Manual corrections were applied to model-generated labels, and the corrected labels were incorporated into subsequent training rounds. After five iterations of refinement, the final segmentation masks were merged with nucleus and plasma membrane labels to create a final mask for visualization. Final mask rendering was performed with ITKSnap ^27^.

## Acknowledgments

This work was supported by the U.S. Department of Energy (DOE), Office of Biological and Environmental Research (KP1601011) and Brookhaven National Laboratory LDRD 24-051. The work used the Laboratory for Biomolecular Structure (LBMS), which is supported by the U.S. Department of Energy, Office of Science, Office of Biological and Environmental Research (KP1607011).

## Author contributions

X. Z. and Q.L. designed the study and experiments. X. Z. and Z. L. performed the experiments. X.Z, Z.L, J. K, L.W. and Q.L. analyzed the data. X. Z. and Q.L. wrote the manuscript with help from other coauthors.

Competing interests: Authors declare no competing interests.

Correspondence and requests for materials should be addressed to Q.L.

## Data availability

The 3-dimensional cryo-vEM data can be accessed upon reasonable request.

## Code availability

The codes are freely available at https://github.com/xzhang0123/vEM

**Supplementary Movie 1**. Sliced-view movie of the cryo-VEM volume of a sorghum stem protoplast.

## Supplementary information

**Supplementary Fig. 1.**
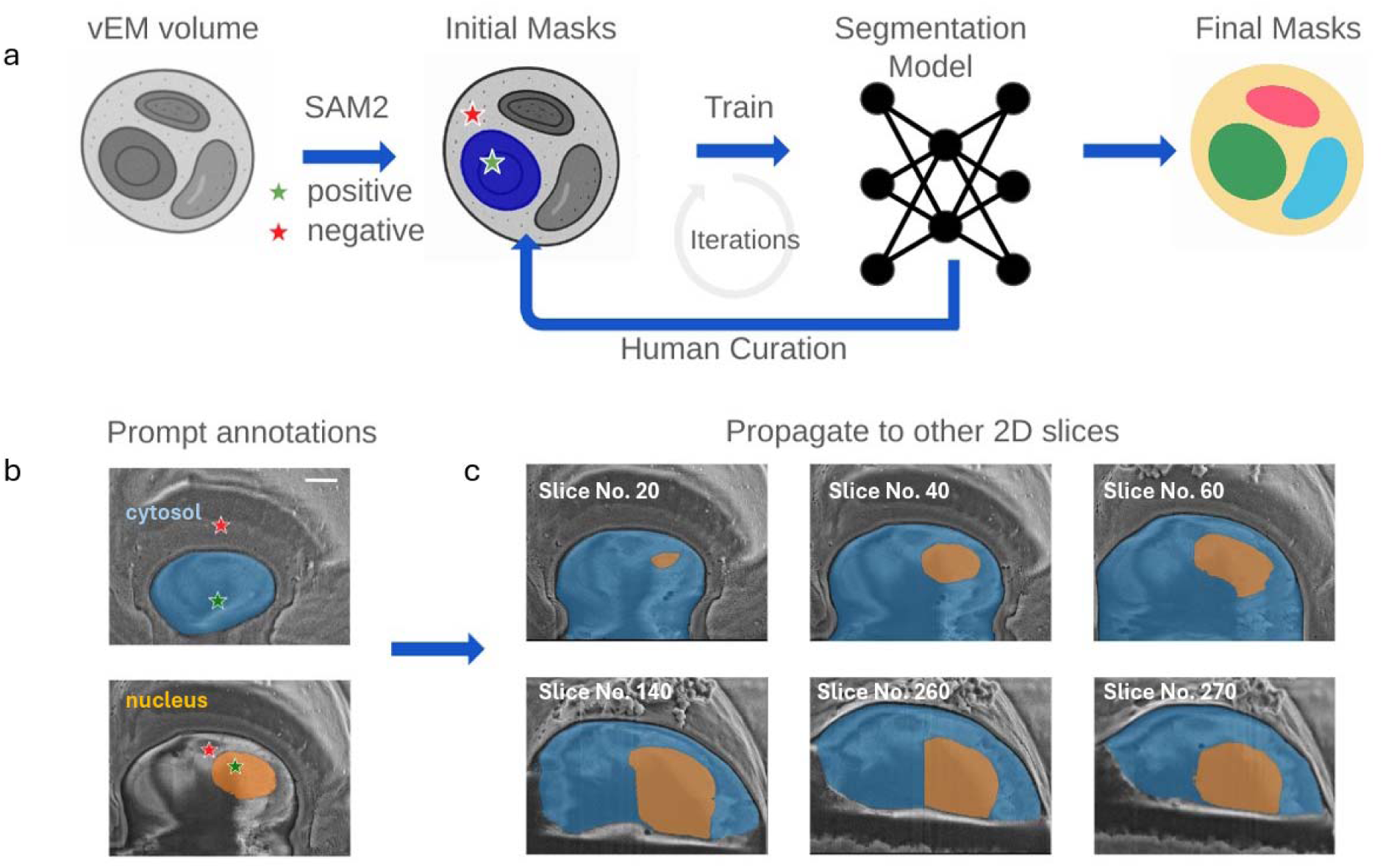
AI-assisted human-in-the-loop segmentation workflow. (a) Overview of the segmentation workflow. An initial mask is generated using the SAM2 model in a Jupyter notebook environment. For each segmentation class, positive (foreground; green star) and negative (background; red star) prompts are manually provided on representative 2D slices The SAM2 model propagates these prompts to generate segmentations on adjacent 2D slices. The resulting 3D mask stack is manually curated and used to train the SwinCell segmentation model. Model predictions were further refined through iterative human curation and retraining. After multiple iterations, this yielded a final segmentation mask suitable for downstream quantitative analysis. (b) Example of user-provided prompts for SAM2. Green stars indicate positive (foreground) prompts, and red stars indicate negative (background) prompts. Here, cytosol (blue) and nucleus (orange) regions are labeled. (c) Mask propagation across slices by SAM2, based on prompts shown in (b).

**Supplementary Fig. 2.**
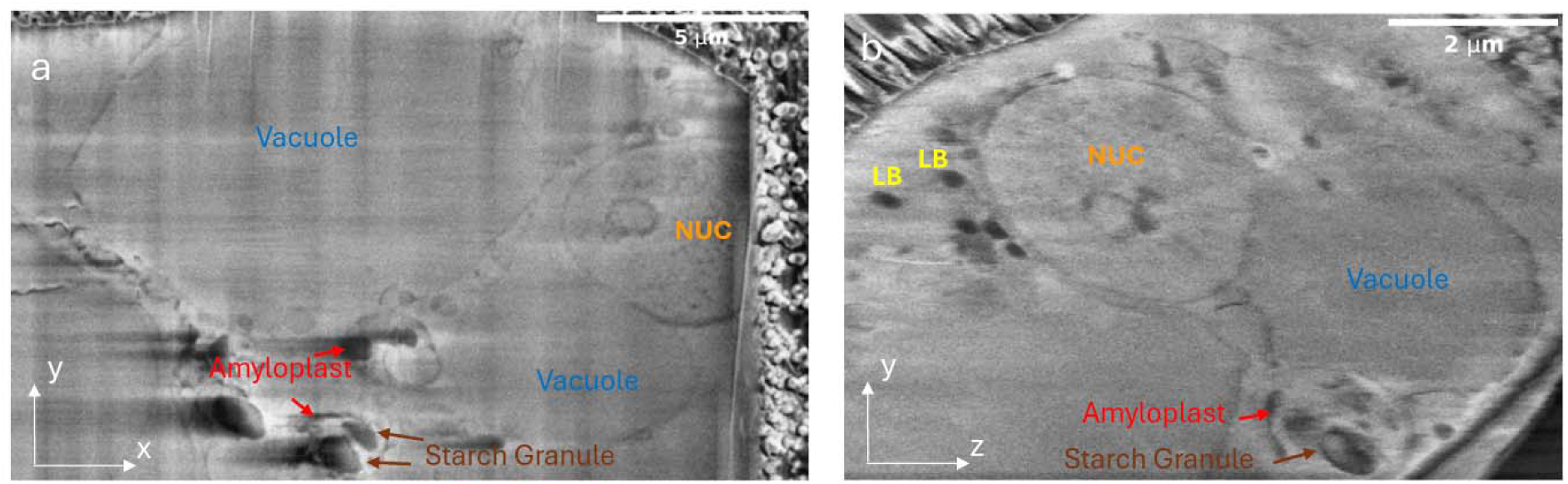
Representative slices extracted from a vitrified sorghum protoplast containing amyloplasts. XY view (a) and YZ view (b) showing key subcellular features, including the nucleus (NUC, orange), vacuole (VA, blue), lipid droplets (LB, yellow), amyloplast-like (red), and starch granule-like (brown).

**Supplementary Fig. 3.**
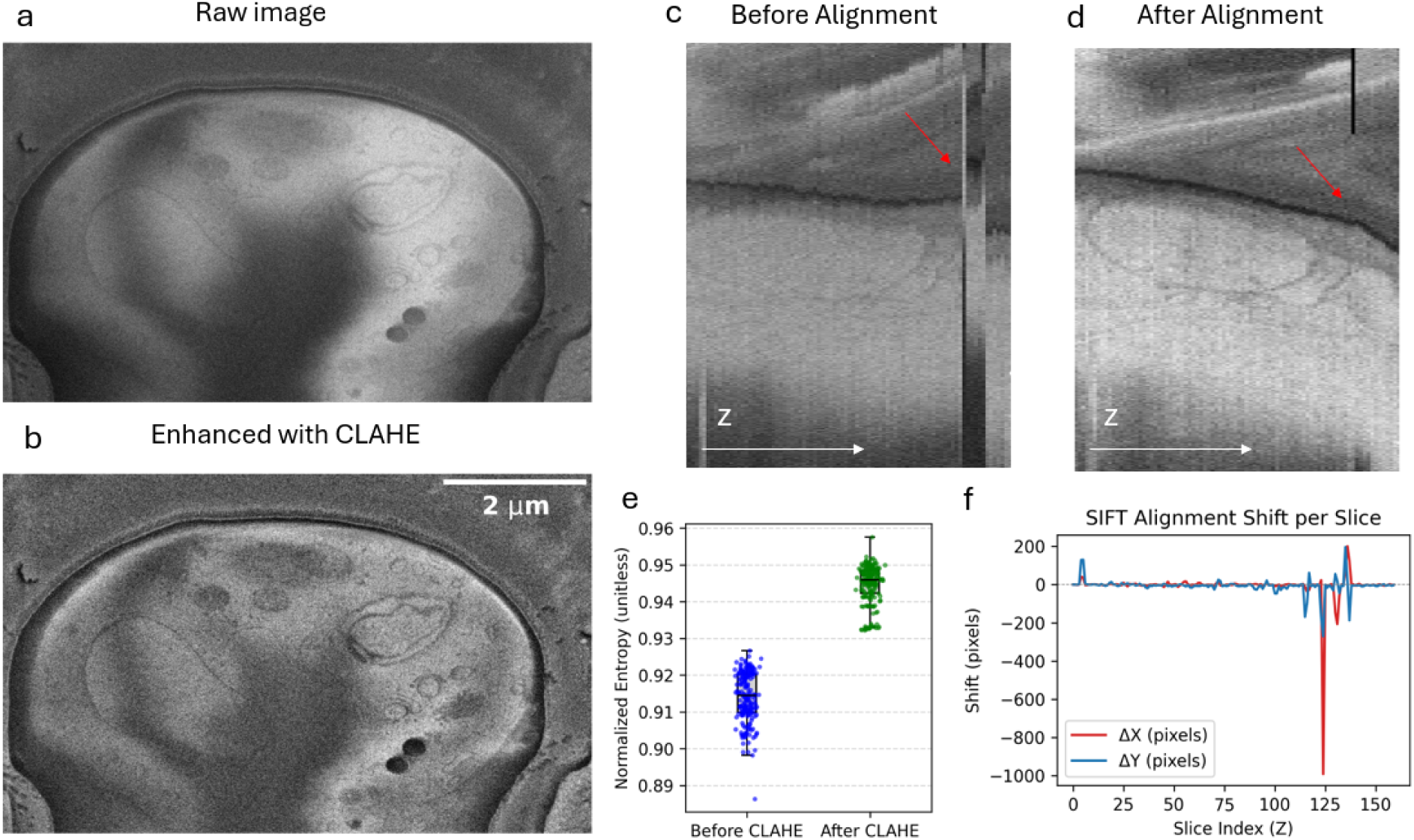
Post-processing of cryo-volume EM (cryo-VEM) data using CLAHE contrast enhancement and SIFT-based alignment. (a,b) Representative XY slices from sorghum protoplast data before (a) and after (b) local contrast enhancement via CLAHE. (c,d) Representative side-view (YZ) slices before (c) and after (d) Scale-Invariant Feature Transform (SIFT) alignment. SIFT improves visualization of subcellular features, including organelle membranes and vesicles. (e) Image contrast of different XY slices measured by Shannon entropy. (f) Measured drifts in X and Y directions determined by the SIFT algorithm; overall, the XY drift exhibits a random pattern.

**Supplementary Fig. 4.**
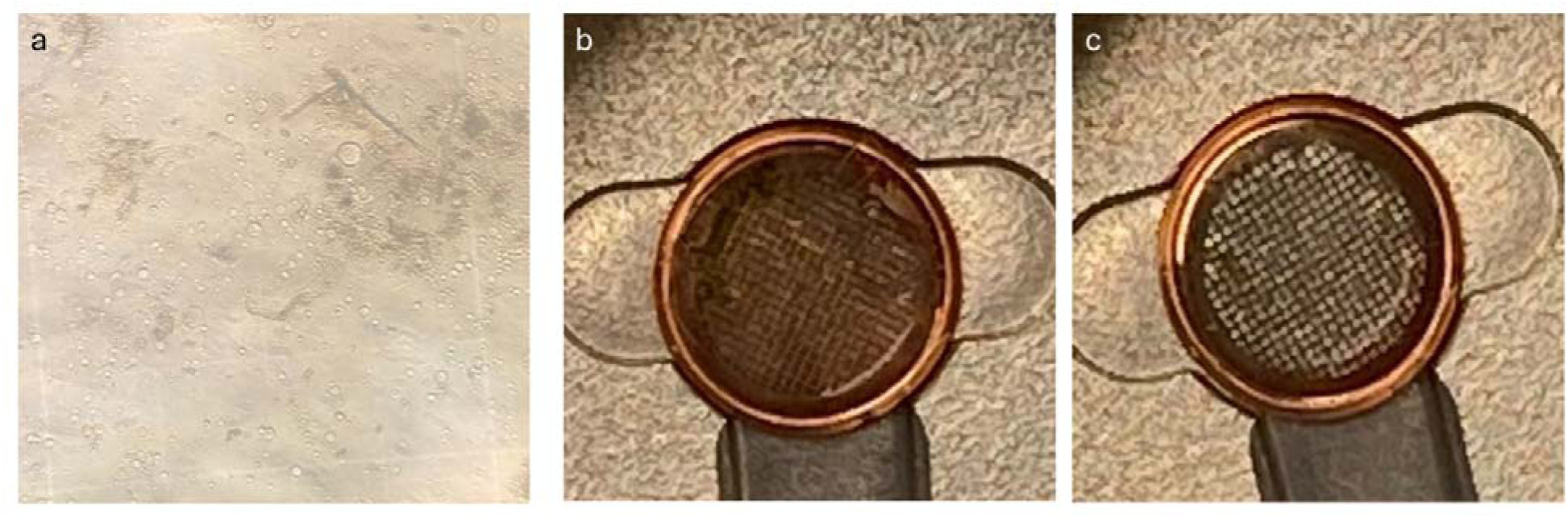
Optimization of protoplast isolation and vitrification conditions. (a) High yield of viable sorghum stem protoplasts obtained with the optimized isolation protocol, visualized using a hemocytometer. (b,c) Assessment of ice thickness after plunge-freezing, evaluated during AutoGrid clipping. Grid transparency and ice uniformity were criteria for suitability in downstream FIB-SEM imaging. (b) Grid with optimal ice thickness. (c) Grid with excessively thick ice, unsuitable for high-resolution imaging.

**Supplementary Fig. 5.**
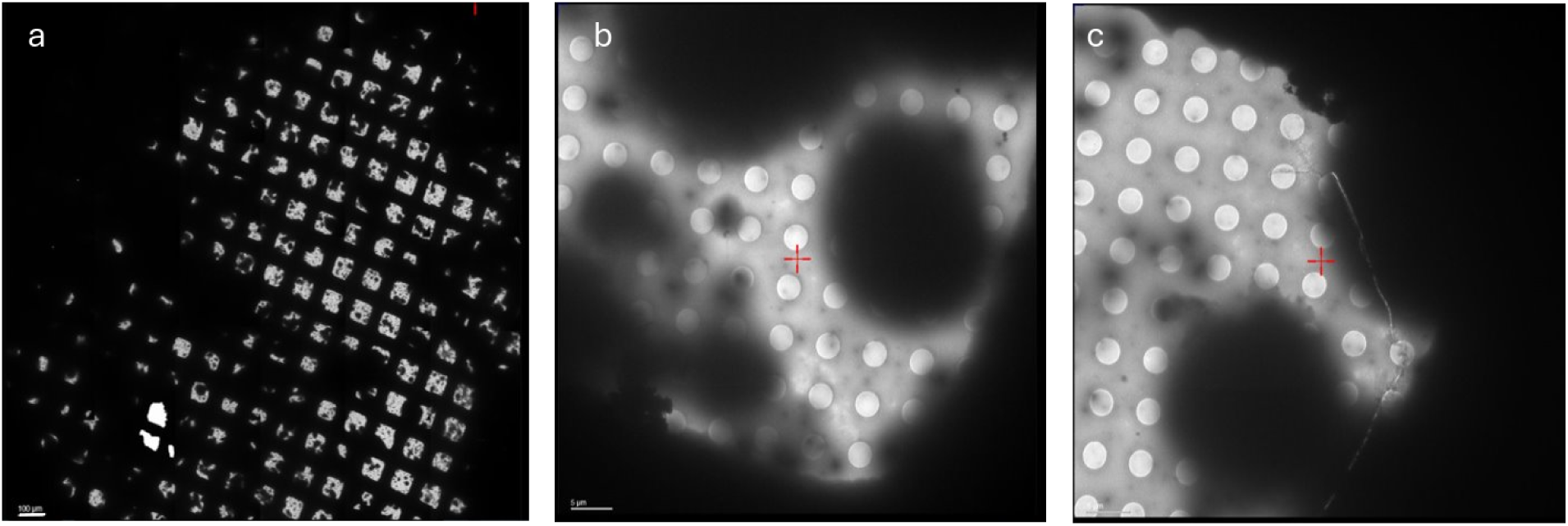
TEM screening of sorghum protoplast distribution and ice thickness at multiple magnifications. Low- (a) and high-magnification (b,c) TEM images were acquired to verify the spatial distribution of sorghum protoplasts on the grid and to assess the ice thickness. Scale bars: (a) 100 μm; (b, c) 5 μm.

**Supplementary Movie 1.** Sliced-view movie of the cryo-VEM volume of a sorghum stem protoplast.

